# An ERP investigation of accented isolated single word processing

**DOI:** 10.1101/2022.07.07.499100

**Authors:** Trisha Thomas, Sendy Caffarra, Clara Martin

## Abstract

Previous studies show that there are differences in native and non-native speech processing (Lev-Ari, 2018). However, less is known about the differences between processing native and dialectal accents. Is dialectal processing more similar to foreign or native speech? To address this, two theories have been proposed. The Perceptual Distance Hypothesis states that the mechanisms underlying dialectal accent processing are attenuated versions of those of foreign (Clarke & Garrett, 2004). Conversely, the Different Processes Hypothesis argues that the mechanisms of foreign and dialectal accent processing are qualitatively different (Floccia et al, 2009). The present study addresses these hypotheses. Electroencephalographic data was recorded from 25 participants who listened to 40 isolated words in different accents. Event-Related Potential mean amplitudes were extracted: P2 [150-250 ms], PMN [250-400 ms] and N400 [400-600 ms]. Support for the Different Processes Hypothesis was found in different time windows. Results show that early processing mechanisms distinguish only between native and non-native speech, with a reduced P2 amplitude for foreign accent processing, supporting the Different Processes Hypothesis. Furthermore, later processing mechanisms show a similar binary difference in the processing of the accents, with a larger PMN negativity elicited in the foreign accent than the others, further supporting the Different Processes Hypothesis. Results contribute to the understanding of single word processing, in which it is uniquely difficult to extract acoustic characteristics from foreign accent, and in which foreign accented speech is associated with the largest cost, as compared to native and dialectal speech, of phonological matching between representations and acoustic input.

**Highlights:**

- Accent affects early speech processing mechanisms at the level of the isolated word
- Acoustic characteristic extraction is more difficult for foreign-accented speech
- Phonological normalization of foreign-accented speech is uniquely difficult
- Accent no longer affects later speech processing mechanisms at the isolated word

## 1. Introduction

It is well documented that unfamiliar accents can lead to difficulties in speech comprehension (Munro and Derwing, 1995; Schmid and Yeni-Komshian, 1999; Anderson-Hsieh and Koehler, 1988; Major et al., 2002). It has furthermore been shown that non-native speech can lead to even greater difficulties (Floccia et al., 2006). As conversations with foreign and regionally accented speakers become increasingly likely, it is important to understand the processing mechanisms involved in these non-standard types of speech comprehension.

Within the realm of accented speech research, much of the research has focused on non-native, or foreign, accented speech processing. However, studying accents in a native, non-native binary does not reflect the true level of complexity of cross-cultural interactions. Often, we may also speak with people who share our native language but are from a different country or region than us. These accent types, called regional or dialectal accents, would have native phoneme variations, unlike foreign accents, and thus it is important to understand the spectrum of accent processing and how processing mechanisms may differ between them. Two hypotheses have been proposed to compare processes underlying foreign and regional accent adaptation.

The Perceptual Distance Hypothesis relies upon the knowledge that accents can be placed on a perceptual scale based on their acoustic distance from native speech, foreign accents being the furthest from native and regional accents falling somewhere in between. According to this hypothesis, regional accent processing mechanisms would be attenuated versions of foreign-accented processing mechanisms (Clark & Garrett, 2004). Behavioral evidence for this hypothesis was reported by Adank et al. (2009), who found that processing speed decreased as accents were less familiar or foreign. Floccia et al. (2006) similarly favored this hypothesis when they found that spoken words were recognized more slowly when produced in an unfamiliar accent, and even more slowly in a foreign accent, as compared to the participants’ native accent.

However, other studies have found data inconsistent with this theory, including additional studies by Floccia (Floccia et al., 2009, Girard et al., 2008). This led to the proposal of the Different Processes Hypothesis, which posits that there are qualitative differences in the processing mechanisms recruited for regional and foreign speech normalization (Floccia et al., 2009). Studies such as Floccia et al. (2009) and Girard, Floccia & Goslin (2008), mentioned above, have supported this hypothesis by showing that French speaking children (Girard et al., 2008) and English speaking children (Floccia et al., 2009) are not able to reliably distinguish between regional and native accents while they are able to distinguish between foreign and native accents, even when accents strengths between the non-native accents are equated, suggesting differences in the linguistic characteristics of the respective speech signals and required normalization mechanisms. Evans and Taylor (2010) also supported the idea of differential processing for foreign and regional accents by showing easier adaptation to foreign than regional accents, despite slower and more error-prone responses with the foreign accents, in a comprehension-in-noise task.

Along with behavioral techniques of studying accented speech comprehension, electrophysiological methods have been increasingly employed due to their ability to provide valuable time-sensitive information. Within this technique, several event-related potentials (ERPs) have been found to be related to various processes during spoken word recognition.

One such component of interest is the P2 or P200 component, which is a positive-deflecting, anterior waveform peaking at around 200 milliseconds (ms) post-stimulus and often distributed around the centro-frontal area of the scalp. This component is associated with the extraction of acoustic features in speech (Reinke et al., 2003). The amplitude of the P2 increases with ease of extraction and is smaller in situations where acoustic extraction is more difficult. In the context of accented speech, a reduction in the amplitude of the P2 has been observed with foreign-accented speech reflecting an increased difficulty in extracting phonetic information (Romero-Rivas et al., 2015). This study did not include a dialectal accent contrast so it is difficult to say which hypothesis their findings support. Another study that included both a dialectal and foreign accent, aimed to investigate the effects of in-group membership and confidence on trust (Jiang et al, 2020). They found a reduction in the P2 only for the dialectal accented condition. However, they note in their discussion that while the native and foreign accents used (Canadian accent and Quebec French accent) were highly familiar to the participants due to the geographical region of the data collection, the dialectal accent (Australian) was much less familiar, which could have influenced the results (Jiang et al., 2020).

Another component of interest is the Phonological Mismatch Negativity (PMN), a negative-going waveform that generally peaks between 250 and 300 ms post-stimulus onset and is usually distributed along a centro-frontal line. The PMN is related to normalization between acoustic input and phonological representations (Newman & Connolly, 2009; Goslin et al., 2012). This component is thought to reflect pre-lexical level processing (Connolly et al., 2001; Newman et al., 2003). Phonological incongruencies, or a mismatch between perceived speech input and a lexical representation, are thought to elicit a larger PMN (Connolly and Phillips, 1994). Because of this finding, the PMN has been described as representing a ‘goodness-of-fit’ marker between phoneme representations and spoken input (Newman & Connolly, 2009). The interpretation of the PMN as a ‘goodness of fit’ marker, is in line with the Perceptual Distance Hypothesis because it supports the idea that the PMN is less pronounced as speech input gets close to native pronunciations (e.g., dialectal or highly proficient non-native speech). However, other recent evidence has argued that the PMN may be better characterized as an index of expectation, rather than an index of phonological mapping, due to their findings that accent experience modulated PMN effects possibly due to experienced participants’ adjusting expectations for non-native speech (Porretta et al., 2017). Other studies such as Goslin et al. (2012) and Sumner & Tilsen (2011) have supported the Different Processes Hypothesis by providing evidence that regional and foreign accents recruit different normalization mechanisms. For example, Goslin et al. (2012) found that the amplitude of the negative-going PMN for regional accents was significantly greater than that of the native accent while the PMN for foreign accents was significantly smaller. This led them to conclude that different strategies were employed to process regional and foreign accents at early processing stages (200-350ms).

Finally, the N4 or N400, a negative-deflecting waveform that usually peaks around 400 ms post-stimulus with a widespread distribution that generally reaches maximum negativity at centroparietal sites, is also of interest when studying spoken word recognition. It has been associated with auditory semantic processing (Bentin et al., 1993; Kutas and Hillyard, 1980) and lexico-semantic integration (Kutas et al., 1987). There have been mixed results on the effects of accent on the N400. Hanulikova et al. (2012) observed a similar N400 effect during listening to both native and foreign-accented spoken sentences with semantic violations (cf. Gosselin et al. 2021). While Goslin et al. (2012) found a reduced N400 effect in correctly spoken sentences during foreign accent listening as opposed to regional or native listening. Goslin et al. (2012) thus argued that participants employed top-down resources to successfully perform word recognition when comprehension was degraded due to foreign accent (relative to dialectal and native). They argued that their results provided further evidence for the Different Processes Hypothesis due to the ability to successfully normalize regional accents during pre-lexical processing while foreign accents could not be normalized and thus continued to have an effect at lexical access and integration stages.

While there is strong evidence that foreign accent processing is different from native accent processing, it is still not clear how dialectal accents are processed nor the relationship between foreign, native and dialectal accent processing mechanisms. Previous studies have contributed to the plausibility of both previously introduced hypotheses, and importantly for our purposes, previous ERP experiments exploring the question have used sentences as material. More studies are needed to elucidate whether foreign accent is processed uniquely from all types of native speech (both native and dialectal accents) or whether dialectal accent is treated differently from native accent, despite both being native speech variations. Exploring accent processing in these three accent types through isolated word listening is useful to clarify accent processing mechanisms without the influence of sentence context, which adds complexity through potential prediction mechanisms and top-down effects. Such a low-level analysis at a small speech unit is convenient to provide a clear, simple picture to advance the debate on the Perceptual Distance and Different Processes Hypotheses. Thus, this will be the focus of the present study.

### 1.1. The Present Study

The present study aims to investigate the processing mechanisms of accented speech using electrophysiological methods. We hope to clarify the processing of foreign, dialectal and native accented speech through the lens of previously proposed hypotheses about accent processing. Specifically, we aim to reveal more about how we process dialectal accent and whether it is treated more similarly to a foreign accent or a native accent during isolated word recognition. ERPs were used because their rich temporal resolution was especially appropriate to study the time course of isolated-word processing mechanisms.

If results support the Perceptual Distance Hypothesis, dialectal accent processing should be more similar to foreign accent processing, although with attenuated effects. Thus, according to the Perceptual Distance Hypothesis, we should observe a gradient in the ERP responses in one or more ERP components that are crucial for speech recognition.

In contrast, a situation where the brain is mainly sensitive to a native/non-native distinction would be in line with the Different Processes Hypothesis, meaning that foreign-accented speech will be treated differently from both dialectal and native accent processing.

## 2. Materials & Methods

### 2.1. Participants

Thirty Spanish natives^1^ participated in the study. Five participants were excluded from further analyses due to excessive artifacts in the electroencephalogram (EEG) recording or due to data collection error (missing behavioral data of one participant). The final sample of participants consisted of 25 females^2^ (mean age: 23 years, SD: 3.54, age range: 19-31 years, Spanish age of acquisition: 0). A post hoc power analysis of the final sample (n=25), using the software G* Power (Faul & Erdfelder, 1992) revealed the power to detect a medium sized effect (f=0.25; cf Cohen, 1977) with alpha at 0.05 was 0.76. All participants lived in the Basque country and considered the Basque-Spanish accent as their native accent. All participants were right handed and had normal or corrected-to-normal hearing and vision. No participant reported a history of neurological disorders. All participants signed an informed consent form before taking part in the study that was approved by the Basque Center on Cognition, Brain and Language ethics committee. They received monetary compensation for their participation.

### 2.2. Materials

Forty Spanish words were extracted from dialogues recorded by 6 female speakers in their thirties with differing accents and presented three times (tot: 120 words per accent). The words were recorded in a Basque Spanish native accent, a Cuban Spanish accent and an Italian Spanish accent by two females for each accent (i.e., hereafter native, dialectal and foreign accent, respectively). The native speakers were born and lived in Spain (Basque country). The dialectal and foreign speakers were chosen for their strong accents and high level of Spanish (in the case of the Italian speakers). Overall, the recordings did not differ in duration (ms) across accents [foreign: 519.7, SD:128.7; native: 505.7, SD: 137.2; dialectal: 519.9, SD:131.6; one way ANOVA: (F(2,117)=0.15, p =0.86)]. Accent strength ratings collected from a separate normative study consisting of eleven participants (average age =23, SD=10) who completed a short online survey where they listened to clips of each accent and rated the accent strength from 1 (mild accent) to 5 (strong accent), showed a clear effect of accent (one way ANOVA: F(2,10)=6.01, p=0.009). Follow up analyses of the clip ratings corrected with the Fisher’s Least Significant Difference (LSD) showed that the dialectal accent was marginally significantly different than the native accent (t(10)=2.13, p=.06), the foreign accent was significantly different from the native (t(10)=3.16,p=.00) and the dialectal and foreign accents were not rated significantly different from each other in terms of strength (t(10)=1.15, p=.28).

### 2.3. Procedure

Participants were seated in a sound-attenuated room in front of a computer screen and were asked to listen to words and occasionally repeat the last word they heard when instructed to by writing on the screen (to maintain listeners’ attention). Each trial began with the symbol *.* at the center of the screen, which was followed by a 500-msec fixation cross. Then a word was presented through speakers while a fixation cross was displayed on the screen. Thirty-three percent of the words were followed by the Spanish word for repeat (‘repite’) displayed on the screen to indicate that the participant should repeat the word that they heard. To minimize artifacts in the EEG recording during the presentation of the auditory stimuli, the participants were asked to blink only when the symbol *.* was presented on the screen. The experimental session lasted about an hour and was divided into 9 blocks. We adopted a blocked design where each accent type was presented in 3 consecutive blocks. The accent presentation order was counterbalanced across participants. Within each block, 40 words of the same accent type were presented in a randomized order (20 words for each speaker).

### 2.4. EEG Data Recording and ERP Analyses

The EEG signal was recorded from 27 channels placed in an elastic cap: Fp1, Fp2, F7, F8, F3, F4, FC5, FC6, FC1, FC2, T7, T8, C3, C4, CP5, CP6, CP1, CP2, P3, P4, P7, P8, O1, O2, Fz, Cz, Pz (see fig.1). Two external electrodes were placed on the mastoids, two were on the ocular canthi, one above and one below the right eye. All sites were referenced online to the left mastoid. Data were recorded and amplified at a sampling rate of 250 Hz. Impedance was kept below 10 KΩ for the external channels and below 5 KΩ for the electrodes on the scalp. EEG data were re-referenced offline to the average activity of the left and right mastoid. A low-pass filter of 30 Hz and a high-pass filter of 0.01 Hz were applied. Vertical and horizontal eye movements were corrected following the Independent Components Analysis (ICA). The fast ICA restricted method was used (1470.3 s). The EEG of each participant was decomposed into independent components and components that explained the highest percentage of the variance in the Veog and Heog channels were identified. Data inspection was then performed on these components to ensure they were accurately accounting for variability related to ocular artifacts. Residual artifacts exceeding ± 100 uV were rejected. On average, 13% of trials were excluded. The number of rejections did not differ across conditions (one way ANOVA: F(2,72)=0.20, p =0.82). For each target word, an epoch of 1200 msec was obtained including a 200 msec pre-stimulus baseline. Average ERPs time-locked to the word onset were computed and baseline corrected for each condition. Statistical analyses were carried out in the following time windows: 150-250, 250-400, and 400-600 msec. The temporal boundaries of each time window were defined based on visual inspection and were also similar to those used in previous ERP studies on auditory word comprehension (Hanulíkova et al., 2012; Rossi, Gugler, Friederici, & Hahne, 2006; Schmidt-Kassow & Kotz, 2008; Van Berkum et al., 2008).

**Fig. 1.**
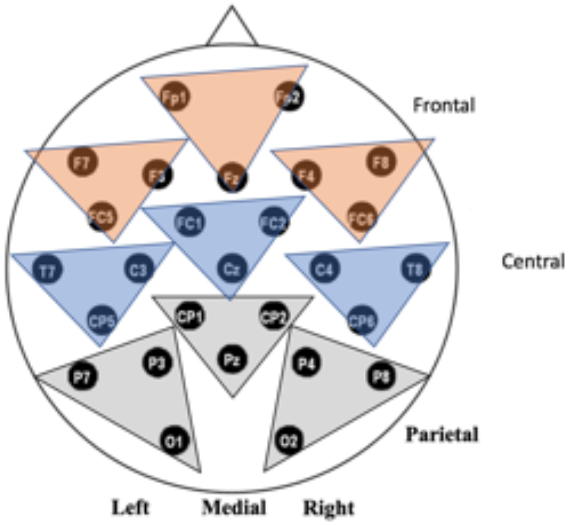
Schematic of electrode montage with topographic organization labeled.

Repeated measures analysis of variance (ANOVA) using conservative degrees of freedom (Greenhouse and Geisser, 1959) was run on the average waveform amplitudes within the time windows of interest. Each ANOVA consisted of accent as a three-level factor (native, dialectal, foreign) and included a topological organization of longitude (frontal, central, parietal) and latitude (left, medial, right). All electrodes were considered in the ANOVA in order to enhance statistical power and directly test the distribution of the effect without theoretical constraints. Each longitude x latitude combination was linearly derived from a combination of three sites: left frontal(F7, F3, FC5), medial frontal(FP1, FP2, Fz), right frontal (F4, F8, FC6), left central (T7, C3, CP5), medial central(FC1, FC2, CZ), right central(C4, T8, CP6), left parietal(P7,P3,O1), medial parietal (CP1,CP2, Pz), right parietal(P4,P8,O2). Finally, t-test post hoc comparisons were conducted and corrected for multiple comparisons using the Benjamini-Hochberg procedure.

## 3. Results

### 3.1. Behavioral Results

Participants showed high accuracy during the online word repetition task meant to test attention and intelligibility (mean overall accuracy: 93.8%, SD:6.8). The word repetition scores did not differ between accents [foreign: 95.3%, SD:10.3; native: 93.6%, SD: 7.6; dialectal: 92.6%, SD:10.0; one way ANOVA (F(2,72)=0.53, p=.59)] showing that the participants could perceive and repeat the words equally well in every accented condition.

### 3.2. ERP results

Average ERPs time-locked to 200 ms pre-word onset in native, dialectal, and foreign accent conditions can be seen in Fig. 2. Topographic distribution for each accented comparison that reached significance was calculated in the 150-250 ms and 250-400 ms time windows and can be seen in Fig. 3 and 5. Plots showing the individual results and distribution of data for each time-window of interest can be seen in Fig. 4, 6, and 7.

**Fig 2.**
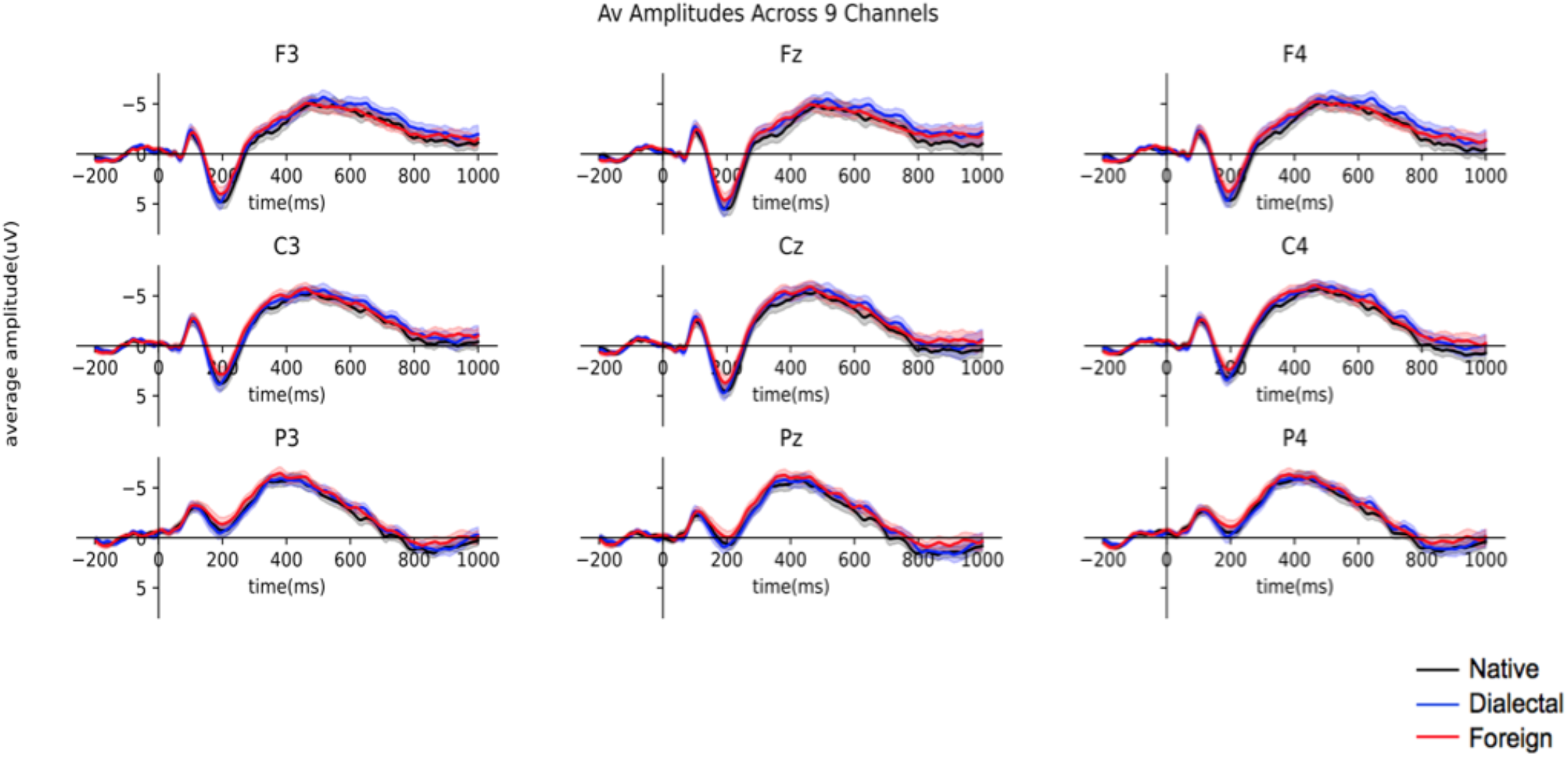
Grand average ERP amplitudes from 200ms pre-word onset until 1000ms after, in different accents. Negativity is plotted up. One standard error is shown.

**Fig. 3.**
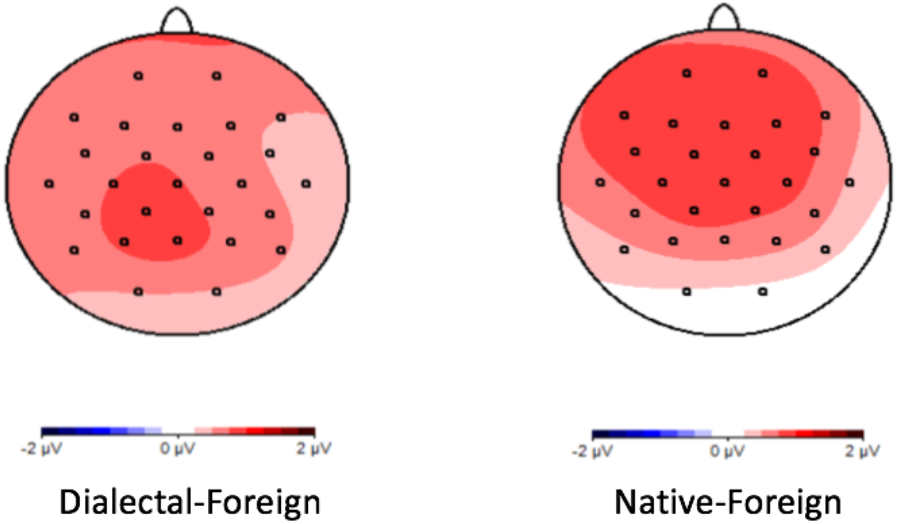
Topographic distribution of voltage difference between conditions with significant differences between 150-250 ms.

**Fig. 4.**
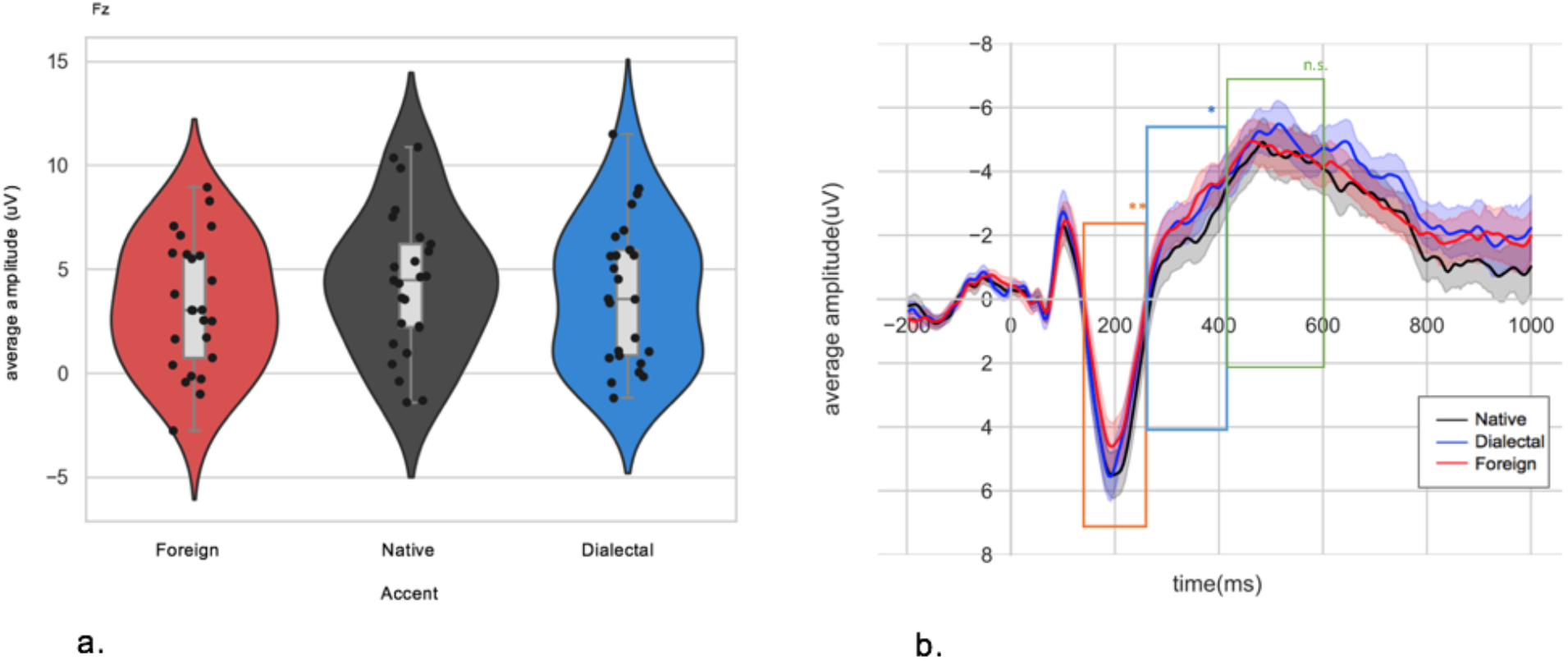
a. Violin plots of the average amplitude of each condition and individual scatterplots with significant differences between 150-250 ms, inner box plot is shown with median, third quartile and first quartile. b. Grand average ERP amplitude from 200 milliseconds prior to word onset (−200) till 1000ms after, in different accents. Negativity is plotted up. The three time-windows of analyses are highlighted (orange for the P2, blue for the PMN, green for the N400). One standard error is shown.

**Fig. 5.**
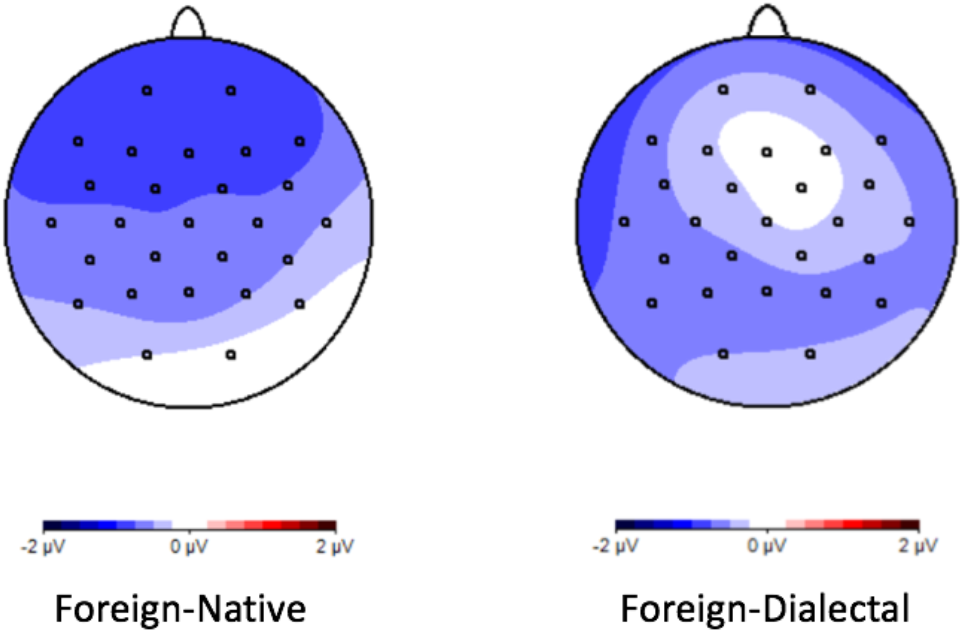
Topographic distribution of voltage difference between conditions with significant differences between 250-400 ms.

**Fig. 6.**
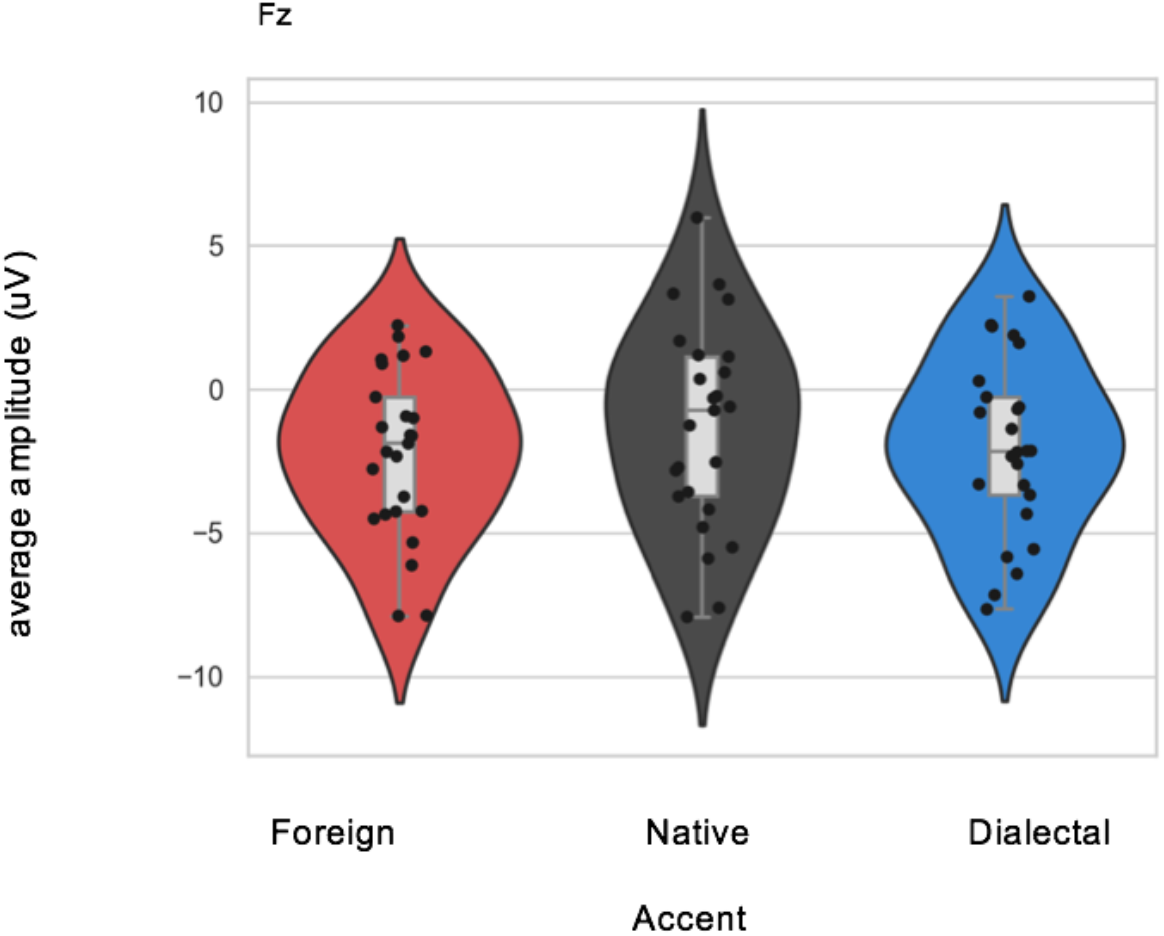
Violin plots of the average amplitude of each condition and individual scatterplots with significant differences between 250-400 ms, inner box plot is shown with median, third quartile and first quartile.

**Fig. 7.**
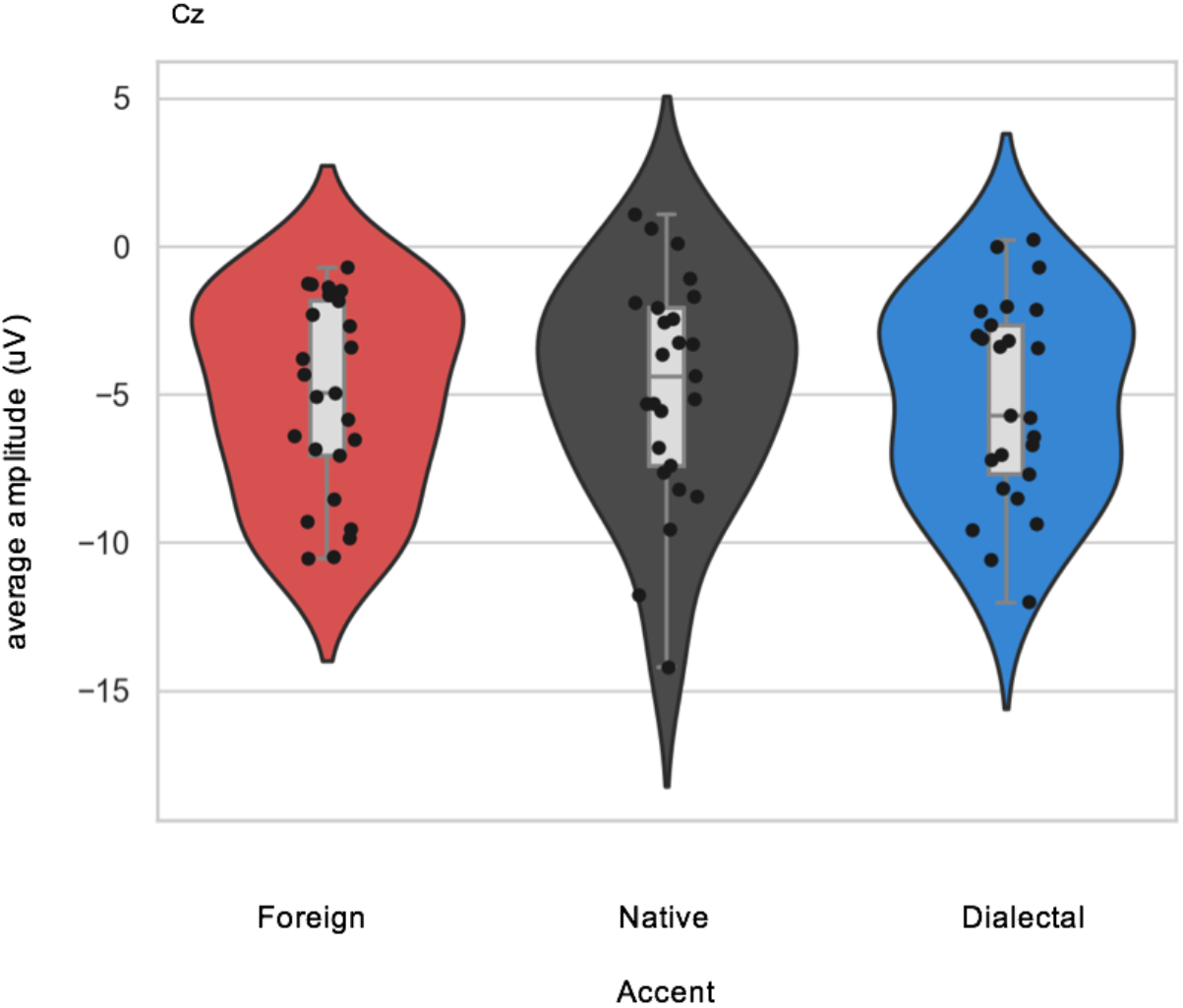
Violin plots of the average amplitude of each condition and individual scatterplots between 400-600 ms, inner box plot is shown with median, third quartile and first quartile.

#### 3.2.1. P2 [150-250 ms]

A main effect of accent was found (F(2,48) = 5.19, p < .05, η^2^=0.34), with a reduced P2 amplitude only for foreign accent (foreign vs native: t(24)=2.92, p<.01, foreign vs dialectal : t(24)=2.88, p<.01, dialectal vs native: t(24)=0.37, p=0.71). The interaction between accent and longitude was also significant (F(4,96) =4.03, p<.05, η^2^=0.33) suggesting that the difference between foreign and native accent was anteriorly distributed (t(24)=3.36, p<.05), while the difference between foreign and dialectal accent was posteriorly distributed (t(24)=3.14, p<.05). All the other contrasts did not reach significance (ps>0.05).

#### 3.2.2. PMN [250-400 ms]

A main effect of accent was found (F(2,48) = 3.43, p < .05, η^2^=0.30), with the highest negativity in foreign accent, followed by dialectal and finally native accent (foreign vs native: t(24)= -3.10, p<.01, foreign vs dialectal : t(24)=-2.67, p<.05). There was no significant difference between the native and dialectal amplitudes (dialectal vs native: t(24)=0.59, p=0.56). The interaction between accent and longitude (F(4,96) =3.52, p<.05, η^2^=0.28) and accent, longitude and latitude (F(8,192)= 2.02, p<.05, η^2^=0.60) was also significant suggesting that the effect of accent was mainly distributed anteriorly for the foreign-native difference (t(24)= -3.59, p<.01) and posteriorly for the foreign-dialectal difference (t(24)= -3.03, p<.01). The posterior effect in the foreign-dialectal contrast was somewhat left-lateralized (t(24)= -3.22, p<.01).

#### 3.2.3. N400 [400-600 ms]

No significant effect of accent was observed (F(2,48) = 0.29, p = .72, η^2^=0.02). Additionally, no significant interactions with accent were found (accent x longitude: F(4,96)=0.60, p = .41, η^2^=0.12; accent x latitude: F(4,96)=0.79, p = .50, η^2^=.18; accent x longitude x latitude: F(8, 192)=0.62, p=.12, η^2^=0.29).

## 4. Discussion

This study aimed to add to the literature on the mechanisms of accented listening comprehension by attempting to advance the debate on two popular spoken language processing theories: the Different Processes Hypothesis and the Perceptual Distance Hypothesis. We were especially interested in contributing to the literature on disentangling dialectal and foreign accents.

Previous literature has shown processing differences between native and non-native accented speech listening (Lane, 1963, Lev-Ari, 2012, 2017; Munro and Derwing, 1995a, b; vanWijngaarden, 2001) and to a lesser extent native and dialectal variations (Adank et al., 2009; Floccia et al, 2006, 2009; Goslin et al, 2012; Girard, Floccia & Goslin, 2008). These differences have led researchers to debate whether accent processing exists on a perceptual scale based on acoustic nearness to native speech (Perceptual Distance Hypothesis) or in distinct categories (Different Processes Hypothesis).

We examined this issue by looking at the effect of accent on the modulation of the P2, PMN and N400 ERP components while listening to isolated words in three accented conditions (foreign, dialectal and native).

Our results align with the Different Processes Hypothesis supporting the idea that foreign-accented speech, at the isolated word level, may present a unique processing challenge that is not present in dialectal speech processing. It further appears that the processing costs of accented speech are strongest early on during the time course of isolated word listening and seem to get weaker over time, suggested by the lack of an effect of accent after 400 ms.

We found that P2 is modulated by accent, with a reduced P2 for foreign accent. This result is in line with previous findings that early stages of speech comprehension are compromised when processing foreign-accented speech due to the phonological properties of foreign-accented speech often departing from those of the native listener (Lane, 1963; Munro and Derwing 1995a,b; van Wijngaarden, 2001). Our results in the P2 were similarly congruent with the conclusions of Romero-Rivas, et al. (2015), suggesting that the extraction of acoustic features from foreign-accented speech is more difficult than from native. Our results furthered this conclusion by also determining that the extraction of acoustic features from foreign-accented speech was also more difficult than from dialectal, while the extraction of both dialectal and native acoustic features was similarly easy. Therefore, ERP results at the P2 support the Different Processes Hypothesis due to the reduction of the P2 only observable in the foreign-accented condition. This leads us to believe that extracting non-native speech sounds presents a unique challenge that is not triggered by native but non-local speech sounds (i.e. dialectal). This processing effort was not seen for our dialectal condition despite the fact that the Cuban dialectal accent used was considered strongly accented by our normative study participants and phonologically deviated significantly from our native accent of Spanish. This may mean that the ‘coherent deviations’ (see Wells,1982; Goslin et al. 2012) that are present in dialectically accented speech are easily adapted to by listeners and thus do not interfere with the extraction of spectral information and other acoustic features, while foreign deviations make this extraction more difficult.

We also found that the PMN is modulated by accent, with the largest mis-match negativity elicited from foreign accent, reflecting increased resources required by normalization processes. This finding is partially in line with previous conclusions drawn about the PMN that support the Different Processes Hypothesis, such as Goslin et al. (2012), who compared the ERP’s associated with the perception of fully formed sentences spoken in different accents. They found that the amplitude of the PMN elicited by the foreign accent and that by the dialectal accent were significantly different from each other. However, our findings diverged from theirs in the directions of amplitudes found and a lack of significant difference in the native-dialectal comparison. Whereas Goslin et al. (2012) found that the amplitude of the PMN for dialectal accent was significantly greater than that of the native and the foreign was significantly smaller than that of the native, we found a gradient in amplitudes moving from native to foreign, with foreign accent eliciting the greatest negativity. Thus, our findings are more in line with the idea of the PMN as a ‘goodness of fit’ marker. Although numerically we did observe a gradient in the PMN averages of our three accented conditions, the difference between the PMN elicited by native and dialectal accents was not significant thus this study did not provide evidence supporting the Perceptual Distance Hypothesis. However, because the trend of the amplitudes in the PMN was consistent with the Perceptual Distance Hypothesis, we conducted an exploratory Bayesian analysis in order to elucidate a clearer picture of pre-lexical accent processing mechanisms and verify the reliability of the null results. This analysis revealed that there was a strong effect between the foreign and other two accents (BF= 1.6e+9; BF =1.3+5) while there was moderate evidence for the null hypothesis (i.e., lack of difference between native and dialectal accent) in the PMN (BF = 0.42), adding further credibility to support of the Different Processes Hypothesis.

Finally, by the 400-600 ms epoch, we no longer observed any significant effect of accent. This is, again, partially in line with Goslin et al. (2012), who in the 350-600 ms epoch no longer observed a difference between native and dialectal accent conditions but diverged from their finding of a significant difference in the foreign accent condition. While their study suggested that “non-native phonological variations could not be fully normalized in pre-lexical processing levels, and so had a continued effect at lexical access and integration stages” (pp.102), our results suggest that in isolated word listening, participants are able to normalize foreign accent at the pre-lexical processing level and thus the accent no longer affects the lexical access stage. These differences are most likely due to using isolated words instead of sentences and may indicate flexibility of this processing mechanism.

The results of this study have suggested that non-native accent affects early stages of single word processing. They have further suggested that these effects are uniquely seen for non-native accents rather than also native variations. While our results suggest that at the isolated word level the brain is operating on a native/non-native binary distinction, the question remains about how general this apparently binary mechanism is. Is it truly a generalized cognitive mechanism or perhaps the flexibility of processing is influenced by external factors? One factor that might be relevant is the acoustic distance among different accents. The acoustic features of regional accents might be less salient than those of foreign accents. Additional research can clarify the role of acoustic properties in accent speech perception of isolated words. Future studies can try to generalize these results to different types of participant profiles, based on gender, language, exposure to accents, or other factors of interest. In addition, would we observe a dichotomic treatment of native/non-native accents in situations involving much more top-down processing, such as continuous speech? Future studies using continuous speech, where contextual information is more important, may help to clarify the role of this mechanism. The impact of additional factors, such as acoustic distance and top-down mechanisms, could reconcile differences seen in our study and other studies conducted using sentences that contain additional suprasegmental (e.g. prosody stress, tone) information.

## 6. Conclusion

Our results support the idea that, even during mostly passive isolated-word listening, indexical features, such as accent, affect our processing at early stages. Furthermore, our findings support that foreign accents are processed differently from dialectal or native accents, requiring more effort to extract acoustic characteristics and to normalize them into expected phonological representations. Despite this, it appears that we are able to successfully normalize non-native speech and thus by later processing stages accent no longer affects our processing or ability to integrate speech sounds into lexical information.

## Acknowledgments

This work was supported by the Basque Government [BERC 2018-2021 program]; the Spanish State Research Agency [BCBL Severo Ochoa excellence accreditation SEV-2015-0490]; the H2020 European Research Council [ERC Consolidator Grant ERC-2018-COG-819093 to CDM; Marie Sklodowska-Curie grant 837228 to SC]; the Spanish Ministry of Economy and Competitiveness [PID2020-113926GB-I00 to CDM]; the Basque Government [PIBA18-29 to CDM]; the Italian Ministry of University and Research (Programma giovani ricercatori Rita Levi Montalcini) to SC; and the Programa Predoctoral de Formación de Personal Investigador No Doctor del Departamento de Educación del Gobierno Vasco (PRE_2021_2_0006 to TT).

## Data availability statement

Code and data to reproduce the reported findings are available at [INSERT URL UPON PUBLICATION]

## Author contributions

All authors contributed to the conceptualization of the study. SC and CDM performed the data collection. TT performed the data analysis and carried out the initial manuscript preparation. TT, CDM, and SC collaborated on the final version of the manuscript.

## Ethics approval statement

The study was approved by the Basque Center on Cognition, Brain and Language Ethics Committee.

## Figure Legend

Fig. 1. Schematic of electrode montage with topographic organization labeled.

Fig 2. Grand average ERP amplitude from 200ms pre-word onset until 1000ms after, in different accents. Negativity is plotted up. One standard error is shown.

Fig. 3. Topographic distribution of voltage difference between conditions with significant differences between 150-250 ms.

Fig. 4

a. Violin plots of the average amplitude of each condition and individual scatterplots with significant differences between 150-250 ms, inner box plot is shown with median, third quartile and first quartile.

b. Grand average ERP amplitude from 200 milliseconds prior to word onset (−200) till 1000ms after, in different accents. Negativity is plotted up. The three time-windows of analyses are highlighted (orange for the P2, blue for the PMN, green for the N400). One standard error is shown.

Fig. 5 Topographic distribution of voltage difference between conditions with significant differences between 250-400 ms.

Fig. 6. Violin plots of the average amplitude of each condition and individual scatterplots with significant differences between 250-400 ms, inner box plot is shown with median, third quartile and first quartile.

Fig. 7. Violin plots of the average amplitude of each condition and individual scatterplots with significant differences between 400-600 ms, inner box plot is shown with median, third quartile and first quartile.

## Appendix

|  |  |  |
| --- | --- | --- |
| Afuera | Marquesa | Inocencia |
| Amor | Señoritas | Pausa |
| Azucar | Felicidad | Temores |
| Bajo | Masculinos | Jardineros |
| Cabeza | Sirvientas | Princesas |
| Corriendo | Feliz | Terrible |
| Cotilla | Mejor | Locura |
| Cuento | Sueño | Problemas |
| Deliciosa | Futbol | Tonterias |
| Descalzo | Nosotras | Luz |
| Dinero | Tacita | Ropa |
| Manias | Gente | Vieja |
| Señores | Pañuelos |  |
| Envidia | Talones |  |

## Footnotes

1 Due to their geographical location in the Basque Country, all participants were also fluent in Basque as early L2

2 Only females were recruited for this study in order to avoid cross-gender listening effects (Banaji and Hardin, 1996; Cacciari and Padovani, 2007)

